# Yellow and oxidation-resistant derivatives of a monomeric superfolder GFP

**DOI:** 10.1101/2024.01.25.577236

**Authors:** Fernando M. Valbuena, Adam H. Krahn, Sherzod A. Tokamov, Annie C. Greene, Richard G. Fehon, Benjamin S. Glick

## Abstract

Fluorescent proteins (FPs) are essential tools in biology. The utility of FPs depends on their brightness, photostability, efficient folding, monomeric state, and compatibility with different cellular environments. Despite the proliferation of available FPs, derivatives of the originally identified *Aequorea victoria* GFP often show superior behavior as fusion tags. We recently generated msGFP2, an optimized monomeric superfolder variant of *A. victoria* GFP. Here, we describe two derivatives of msGFP2. The monomeric variant msYFP2 is a yellow superfolder FP with high photostability. The monomeric variant moxGFP2 lacks cysteines but retains significant folding stability, so it works well in the lumen of the secretory pathway. These new FPs are useful for common imaging applications.

## INTRODUCTION

Characterization of the green fluorescent protein (GFP) from the jellyfish *Aequorea victoria* transformed biomedical research by allowing proteins, intracellular compartments, and cells to be tracked by fluorescence microscopy (Tsien, 1998). GFP was mutated to enable excitation by blue instead of ultraviolet light and to create spectral variants such as the yellow fluorescent protein (YFP) and the cyan fluorescent protein (CFP). Among the most popular derivatives of GFP are the “enhanced” variants EGFP and EYFP. Two additional classes of mutations improved the physical properties of GFP derivatives. First, point mutations such as A206K eliminated the weak dimerization that was a property of wild-type GFP to yield monomeric variants such as mEGFP and mEYFP (Zacharias et al., 2002). Second, mutations at multiple locations in the protein increased folding stability to yield variants such as superfolder GFP (Pédelacq et al., 2006). Different types of mutations can be combined to optimize multiple properties. For example, we described a GFP variant termed msGFP2, which has monomerizing and superfolder mutations as well as improved N- and C-terminal peptides that reduce cytotoxicity (Valbuena et al., 2020). msGFP2 is more photostable than the original superfolder GFP, and it is an excellent choice for experiments that involve routine fluorescence imaging.

Since the discovery of *A. victoria* GFP, a variety of other FPs have been identified (Chudakov et al., 2010). Most of those FPs are strongly oligomeric, so mutations have been used to disrupt the oligomerization interfaces. But monomeric derivatives of strongly oligomeric FPs are often less well behaved as fusion tags than monomeric derivatives of *A. victoria* GFP (Costantini et al., 2012; Costantini et al., 2015). Therefore, we undertook further modifications to generate two derivatives of msGFP2.

The first msGFP2 derivative is a yellow fluorescent variant. YFPs such as EYFP and Venus, and their monomeric derivatives mEYFP and mVenus, are widely used (Zacharias et al., 2002; Kremers et al., 2006). Excitation wavelengths for YFPs are less phototoxic than those for GFPs (Wäldchen et al., 2015), and YFPs provide good spectral separation from both cyan and red FPs. A limitation of many YFPs is relatively fast photobleaching, but an mVenus derivative termed mGold was reported to be highly photostable (Lee et al., 2020). Inspired by these advances, we have now created msYFP2, a monomeric superfolder YFP with photostability comparable to that of mGold. When used as a fusion tag in living cells, msYFP2 yielded a substantially more photostable signal than mEYFP and Venus, suggesting that msYFP2 will find utility as an improved YFP.

The second msGFP2 derivative is an oxidation-resistant variant. GFP contains two internal cysteine residues, and in oxidizing environments such as the eukaryotic secretory pathway or the bacterial periplasm, those cysteines often undergo disulfide bond formation to generate aggregated and nonfluorescent FP tags (Feilmeier et al., 2000; Jain et al., 2001). This problem can be alleviated by using superfolder GFP and its derivatives, presumably because stable folding renders the internal cysteines less accessible (Aronson et al., 2011). However, a more reliable way to prevent disulfide bond formation is to mutate both cysteines (Suzuki et al., 2012; Costantini et al., 2015). Such mutations destabilize the FP (Jain et al., 2001), but by starting with superfolder GFP, Snapp and colleagues generated the oxidation-resistant variant oxGFP and its monomeric derivative moxGFP (Costantini et al., 2015). Building on that work, we systematically examined the folding of msGFP2 variants containing different amino acid substitutions of the two cysteines.

The result is moxGFP2, which is no longer a superfolder but retains higher folding stability than EGFP. Functional tests confirmed that moxGFP2 is a well-behaved fusion tag in the lumen of the secretory pathway.

## RESULTS AND DISCUSSION

### Generation of the photostable variant msYFP2

As the first step in engineering an improved YFP, we generated a monomeric superfolder variant. EYFP was modified by introducing the following mutations: the monomerizing mutation A206K (Zacharias et al., 2002), the SYFP2 mutations F46L, F64L, L68V, and S175G (Nagai et al., 2002; Kremers et al., 2006), the folding reporter and superfolder mutations S30R, Y39N, F99T, N105T, Y145F, M153T, V163A, and I171V (Pédelacq et al., 2006; Valbuena et al., 2020), and the N- and C-terminal peptides of msGFP (Valbuena et al., 2020). The resulting variant was termed msYFP. Unfortunately, purified msYFP photobleached quickly, even faster than EYFP (Figure 1A).

**FIGURE 1:**
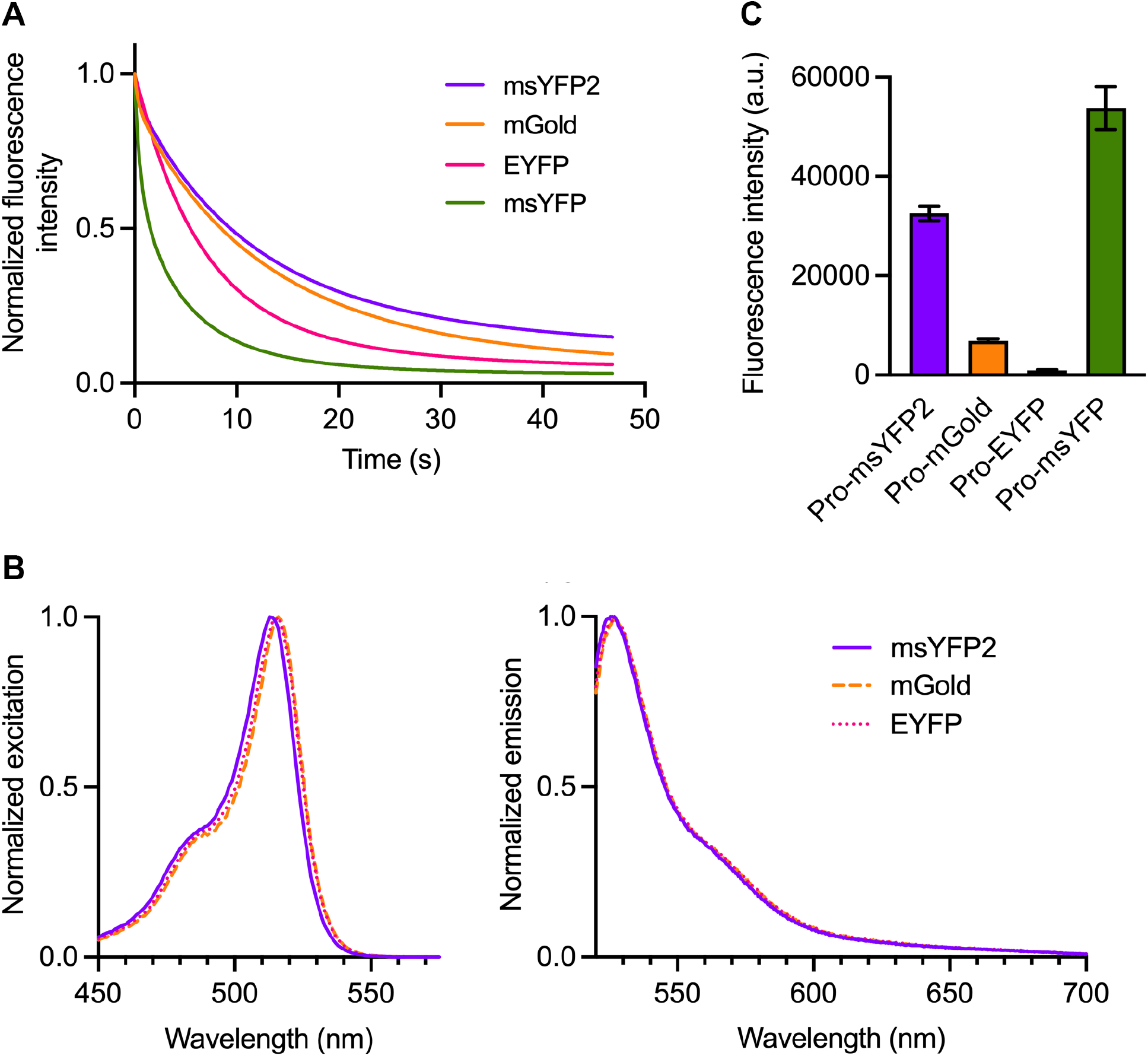
msYFP2 is a photostable superfolder YFP. (A) Photobleaching of purified YFP variants. The indicated YFPs were imaged repeatedly by widefield microscopy. Images were collected every 0.1 s, and the emission signals were quantified. Each curve was normalized to the emission signal from the first image. (B) Spectra of YFP variants. Purified YFPs were diluted to an absorbance at 280 nm of approximately 0.1. Excitation spectra were collected for emission at 527 nm, and emission spectra were collected for excitation at 513 nm. The spectra were normalized to the highest values. (C) Folding stabilities of YFP variants. The indicated YFPs were expressed in *E. coli* as fusions to proinsulin (“Pro”), and the fluorescence signals from the cells were measured. Plotted in arbitrary units (a.u.) are mean and SEM values from three replicates.

We then sought to generate a photostable monomeric superfolder YFP by reverting some of the introduced mutations and by introducing new mutations. During the creation of msGFP2, reversion of the Y145F mutation was crucial for enhancing photostability (Valbuena et al., 2020), but that reversion proved not to be beneficial in the context of msYFP (Figure S1A). By contrast, reversion of the F46L and L68V mutations dramatically enhanced photostability (Figure S1A). A further enhancement was obtained by incorporating the T63S mutation that was described for mGold (Lee et al., 2020). Finally, we introduced the C-terminal peptide of msGFP2 (Valbuena et al., 2020). The resulting variant was termed msYFP2. The absorbance and emission spectra of msYFP2 were very similar to those of EYFP except that the excitation and emission maxima of msYFP2 were blue-shifted by 2-3 nm (Figure 1B). During repeated imaging on a widefield microscope at a fixed excitation intensity, purified msYFP2 showed somewhat greater photostability than mGold and substantially greater photostability than EYFP (Figure 1A). The photophysical properties of msYFP2 are summarized in Table 1.

**TABLE 1:** Photophysical properties of the msGFP2 derivatives. Relative brightness, calculated as the product of the quantum yield and the extinction coefficient, was defined as 1.00 for EYFP and EGFP. *Data taken from Valbuena et al., 2020.

| FP | Absorbance<br>Maximum<br>(nm) | Emission<br>Maximum<br>(nm) | Quantum<br>Yield | Extinction<br>Coefficient<br>(M<sup>-1</sup> cm<sup>-1</sup>) | Relative<br>Brightness |
| --- | --- | --- | --- | --- | --- |
| EYFP | 516 | 527 | 0.67 | 67,100 | (1.00) |
| mGold | 516 | 527 | 0.69 | 114,800 | 1.76 |
| msYFP2 | 513 | 525 | 0.66 | 90,400 | 1.33 |
| EGFP | 489 | 510 | 0.60 | 56,600 | (1.00) |
| moxGFP2 | 488 | 509 | 0.56 | 52,900 | 0.87 |
| msGFP2* | 489 | 509 | 0.59 | 52,200 | 0.91 |

To confirm that msYFP2 has superfolder properties, we fused various YFPs to proinsulin and expressed the fusion constructs in *E. coli* (Valbuena et al., 2020). In the reducing environment of the bacterial cytoplasm, proinsulin cannot undergo disulfide bonding and so it presumably forms aggregates. Non-superfolder FPs that are trapped in those aggregates fail to attain their folded fluorescent states. Indeed, a proinsulin-EYFP fusion was nonfluorescent and a proinsulin-mGold fusion was only weakly fluorescent, whereas a proinsulin-msYFP2 fusion was highly fluorescent (Figure 1C). Those different fluorescence signals reflected different levels of the fusion proteins in the bacterial cells (Figure S1B), likely indicating that the nonfluorescent proinsulin-YFP molecules were degraded. Interestingly, we reported earlier (Valbuena et al., 2020) and confirmed here (see below) that nonfluorescent proinsulin-GFP fusions were not degraded in bacteria. The reason for this difference between YFP and GFP fusions is unclear, but regardless of whether nonfluorescent fusion proteins are degraded, the proinsulin-FP assay is a useful test of folding stability. In sum, msYFP2 is a monomeric FP that combines excellent photostability with high folding stability.

We next confirmed that msYFP2 is photostable *in vivo* by performing three separate tests. In the first test, msYFP2 was expressed in yeast cells as a fusion to the Golgi protein Sec7 (Losev et al., 2006). The Golgi-associated fluorescence was visualized by 4D confocal microscopy, and time-dependent bleaching of the signal was measured (Johnson and Glick, 2019; Valbuena et al., 2020). mEYFP and mGold were used as controls. In accord with the *in vitro* results, the msYFP2 fusion was about as photostable as mGold and much more photostable than mEYFP (Figure 2).

**FIGURE 2:**
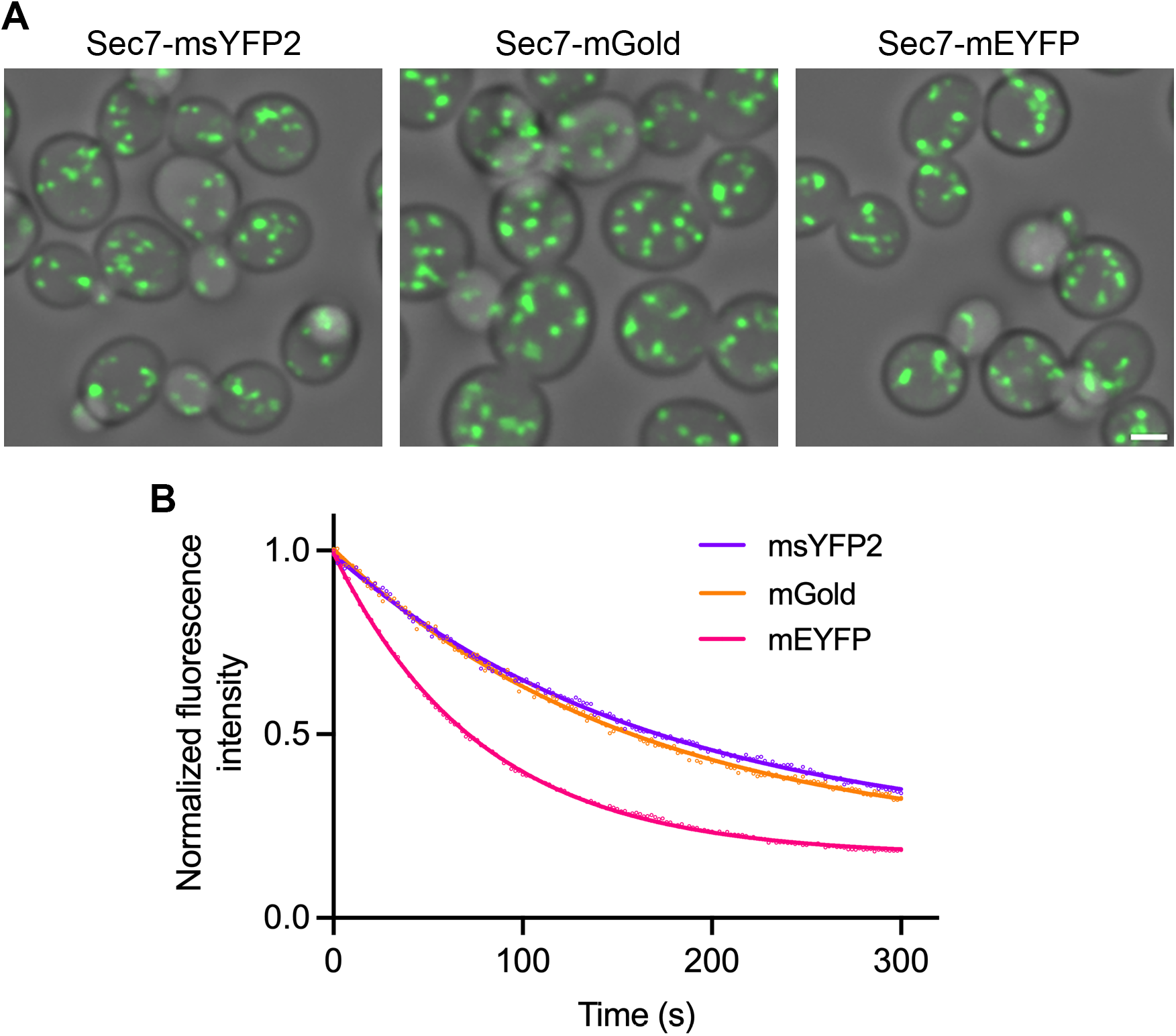
msYFP2 is photostable in live yeast cells. Golgi cisternae were labeled by using gene replacement to fuse the indicated YFPs to Sec7. Confocal z-stacks were captured every 2 s for 5 min and then average projected to create movies. (A) Images from the first frames of representative movies. Scale bar, 2 µm. (B) Plots of averaged fluorescence signals. For each strain, three movies were captured with approximately 10-15 cells in each movie. Averaged fluorescence signals were normalized to the values at time zero, and the data were fit to exponential decay curves. The rate constant for photobleaching of mEYFP was about twice that for the other two YFPs.

In a second test of *in vivo* photostability, we endogenously appended msYFP2 to the *Drosophila* transcriptional co-activator Yorkie (Yki) (Huang et al., 2005) using CRISPR/Cas9 and compared this fusion to a previously generated Yki-Venus fusion (Manning et al., 2020). The Yki-msYFP2 and Yki-Venus fusions were identical except for the YFP portions, and the two constructs were expressed and imaged in parallel. A functional test of the constructs was based on the established finding that when the core pathway kinase Warts (Wts) is depleted, Yki accumulates in nuclei (Figure 3A) (Oh and Irvine, 2008; Xu et al., 2018). Both constructs passed this test, as shown in Figure 3B for Yki-msYFP2 expressed in the *Drosophila* wing imaginal disc. To compare the photostabilities of Yki-msYFP2 and Yki-Venus, we performed consecutive 3D confocal scans of the apical regions of wing disc cells. Some of the tagged Yki molecules presumably diffused between the bleached and unbleached parts of the cells, so this experiment was only semi-quantitative, but the data indicated that Yki-msYFP2 was substantially more photostable than Yki-Venus (Figure 3C).

**FIGURE 3:**
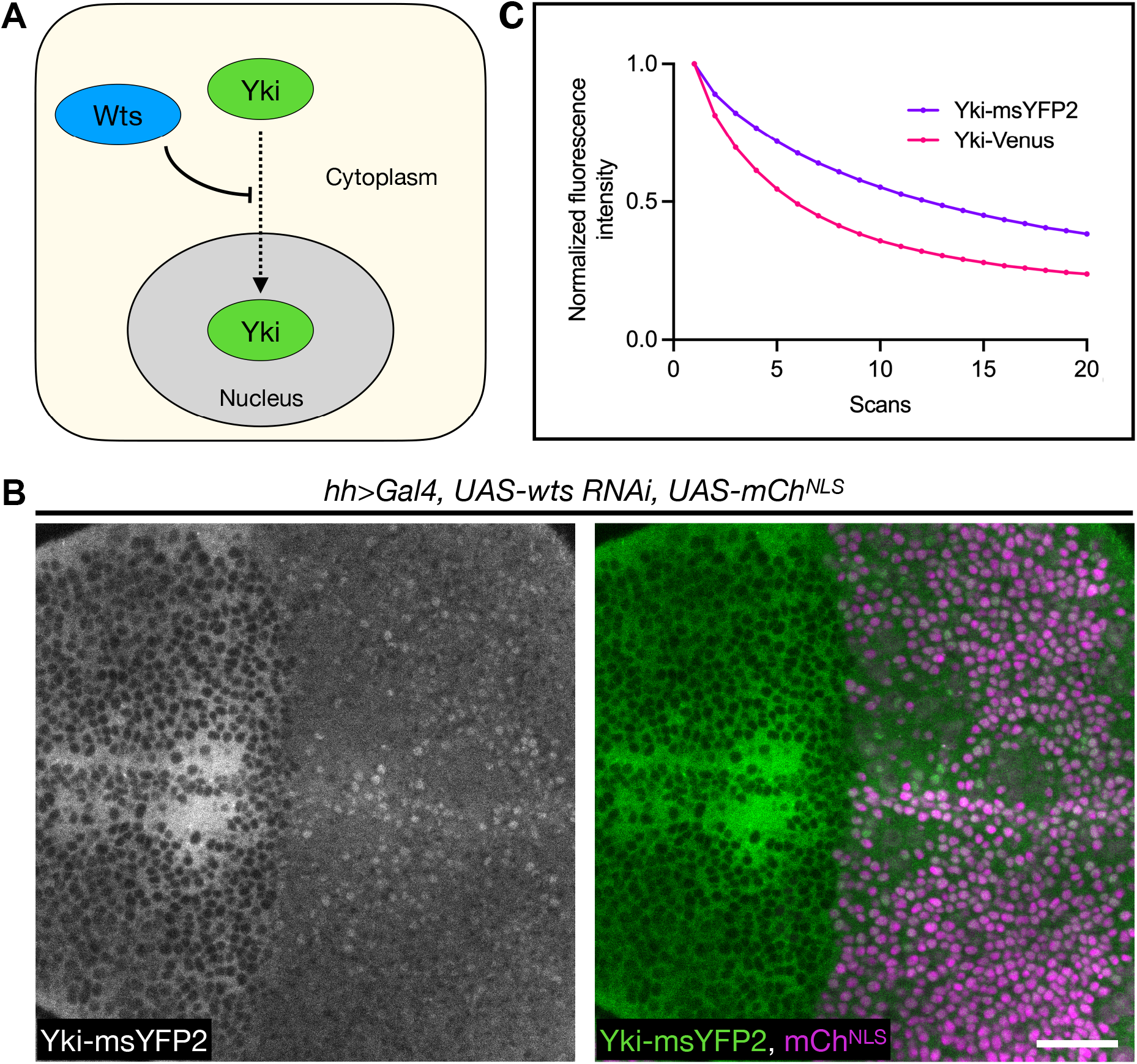
msYFP2 is photostable in *Drosophila* larvae. (A) Simplified diagram of Yki regulation. Wts inhibits the translocation of Yki from the cytoplasm into the nucleus, so in the absence of Wts, Yki enters the nucleus. (B) Depletion of Wts in the posterior compartment of the wing imaginal disc using the *hh>Gal4* driver (mCh^NLS^ positive side) leads to significant nuclear accumulation of Yki-msYFP2 compared to the control anterior compartment (mCh^NLS^ negative side), suggesting that Yki-msYFP2 behaves normally *in vivo*. Scale bar, 20 µm. (C) Plot of Yki-msYFP2 and Yki-Venus photobleaching during scanning confocal imaging of the third instar wing imaginal epithelium. The fluorescence signals were normalized to the values from the first scan.

In a third test of *in vivo* photostability, msYFP2 was compared to mGold by appending these YFPs to the mammalian Golgi protein N-acetylgalactosaminyltransferase 2 (GalNAc-T2) (Storrie et al., 1998). The two constructs were expressed in cultured RPE-1 cells, and Golgi structures were imaged by repeatedly capturing z-stacks. Under these conditions, the msYFP2 and mGold constructs photobleached at similar rates (Figure 4). To measure *in vivo* brightness values for the GalNAc-T2 fusions to msYFP2 and mGold, fluorescence signals were quantified for nocodazole-induced Golgi mini-stacks (Cole et al., 1996). The mGold-labeled mini-stacks were on average about 6% brighter than the msYFP2-labeled mini-stacks. This difference is smaller than would be expected based on the brightness values of purified msYFP2 and mGold (Table 1), perhaps because the superfolder properties of msYFP2 enable it to fold more robustly than mGold in the lumen of the secretory pathway (Costantini et al., 2015). The combined results confirm that msYFP2 is well behaved as a photostable tag for live cell imaging.

**FIGURE 4:**
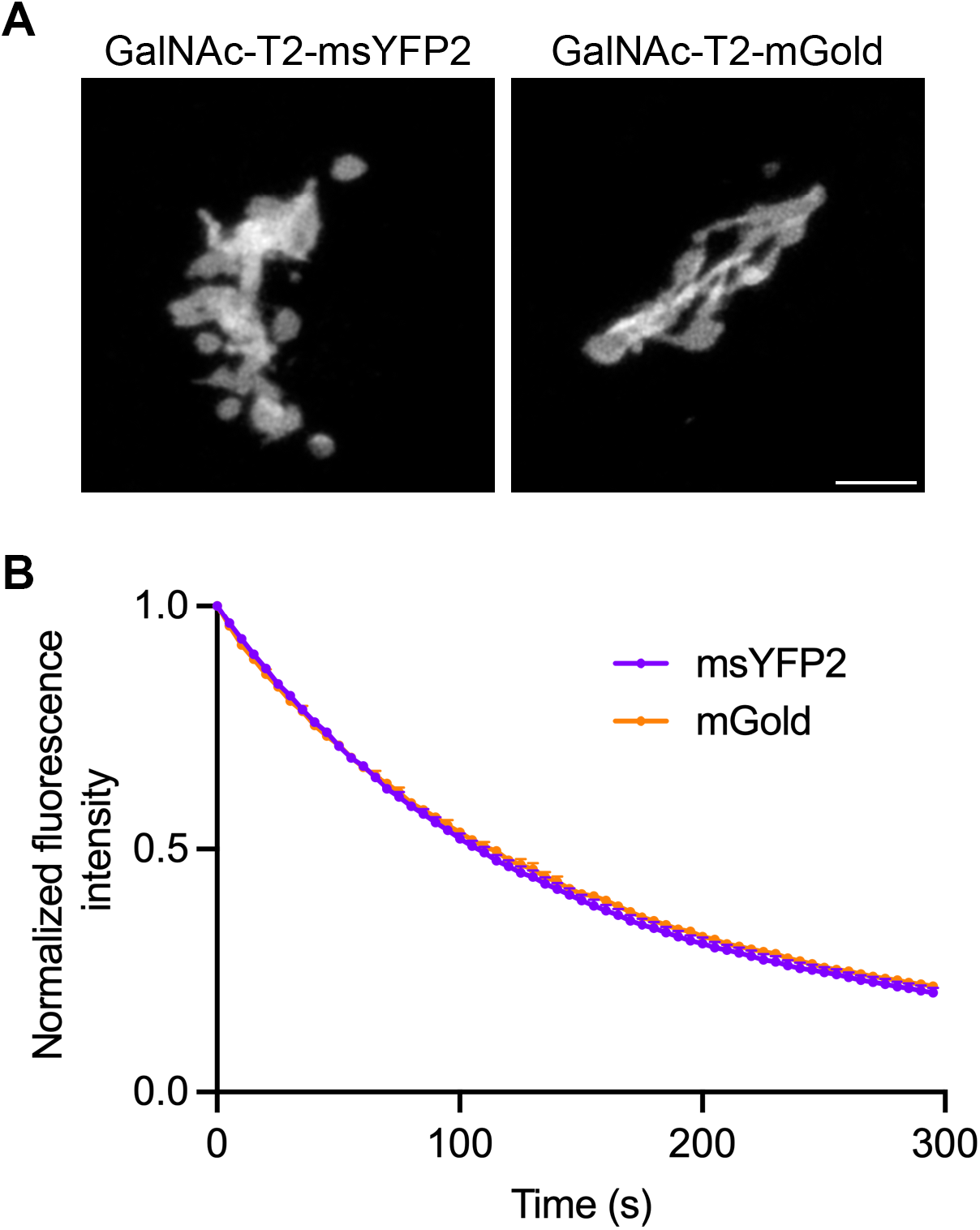
msYFP2 is photostable in cultured mammalian cells. (A) Labeling of the Golgi ribbon in RPE-1 cells by transient expression of constructs encoding msYFP2 or mGold fused to GalNAc-T2. Scale bar, 5 µm. (B) Photostability of msYFP2 and mGold fused to GalNAc-T2 in live RPE-1 cells. Five cells from each sample were imaged by capturing confocal z-stacks every 5 s for 5 min, and the signals from the Golgi structures were quantified. Averaged fluorescence signals were normalized to the values at time zero. The bars represent SEM values.

### Generation of the oxidation-resistant variant moxGFP2

Cys-48 and Cys-70 of GFP can undergo oxidation to form disulfides in some cellular environments, including the ER lumen (Suzuki et al., 2012; Costantini et al., 2015). Mutation of the cysteines destabilizes GFP (Jain et al., 2001), but this effect can be offset by starting with a superfolder variant (Costantini et al., 2015). Our approach was to mutate the cysteines of msGFP2 and then to test the folding stabilities of the resulting constructs using the proinsulin fusion assay. As a guide for choosing alternative residues at positions 48 and 70, we performed a multiple sequence alignment with a diverse set of FPs (Alieva et al., 2008). The amino acid corresponding to Cys-48 is often Val or Ser in other FPs while the amino acid corresponding to Cys-70 shows broader variation. We therefore introduced Val or Ser at position 48 combined with Val, Ser, Thr, or Met at position 70.

After observing that bacterial cultures expressing the C48S/C70T, C48V/C70S, and C48V/C70M mutants exhibited very low fluorescence, we tested the other five mutants by fusing them to proinsulin. As controls, a proinsulin-mEGFP fusion yielded undetectable fluorescence in bacteria whereas a proinsulin-msGFP2 fusion yielded strong fluorescence (Figure 5A). Among the mutants, only the C48S/C70V mutant yielded significant fluorescence when fused to proinsulin (Figure 5A). The fluorescence signal for the C48S/C70V mutant was ∼16-17% as strong as the fluorescence signal for msGFP2 despite similar expression levels (Figure 5A and Figure S2), indicating that the C48S/C70V mutant was no longer a superfolder but retained greater folding stability than mEGFP. The absorption and emission spectra of the C48S/C70V mutant were virtually identical to those of EGFP (Figure 5B). We designated the C48S/C70V mutant moxGFP2. Compared to EGFP, purified moxGFP2 showed slightly higher photostability (Figure 5C) and about 13% lower brightness. The photophysical properties of moxGFP2 are summarized in Table 1.

**FIGURE 5:**
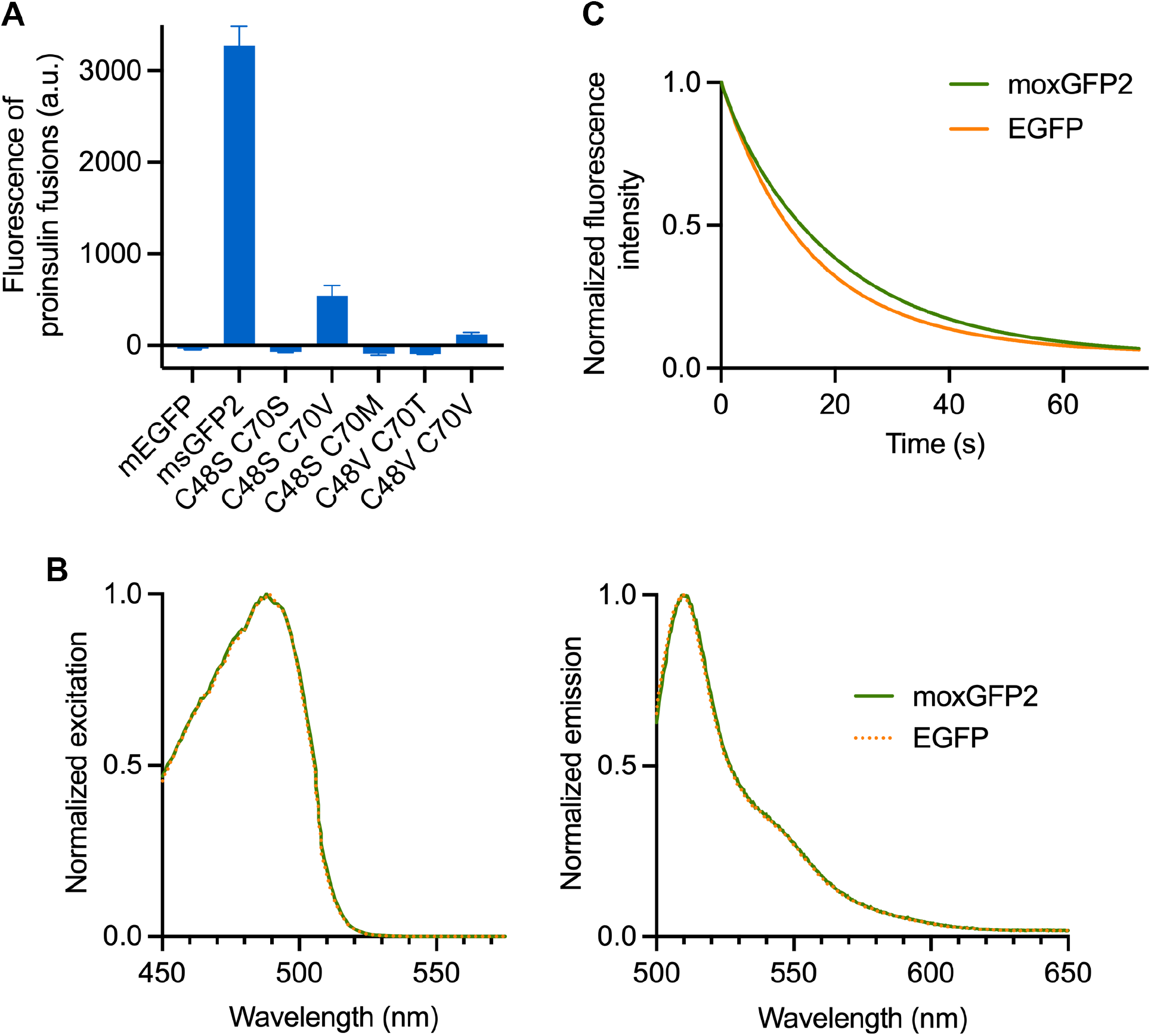
moxGFP2 is a stably folded and photostable oxidation-resistant GFP. (A) Fluorescence of mEGFP, msGFP2, and various msGFP2 mutants when expressed in *E. coli* as fusions to proinsulin. Mean cellular fluorescence is shown in arbitrary units (a.u.), together with SEM values based on nine replicates per sample (three replicates from each of three different colonies). (B) Spectra of moxGFP2 and EGFP. Purified GFPs were diluted to an absorbance at 280 nm of approximately 0.1. Excitation spectra were collected for emission at 507 nm, and emission spectra were collected for excitation at 488 nm. The spectra were normalized to the highest values. (C) Photobleaching of purified GFP variants. moxGFP2 and EGFP were imaged repeatedly by widefield microscopy. Images were collected every 0.1 s, and the emission signals were quantified. Each curve was normalized to the emission signal from the first image.

To test whether moxGFP2 functions in the lumen of the secretory pathway, we performed gene replacements that appended mEGFP, msGFP2, or moxGFP2 to yeast Kar2, which is an Hsp70 protein that resides in the ER. As previously observed (Valbuena et al., 2020), Kar2-mEGFP yielded large fluorescent aggregates and inhibited cell growth, whereas Kar2-msGFP2 yielded a typical ER pattern and did not affect cell growth (Figure 6A,B). Kar2-moxGFP2 behaved like Kar2-msGFP2 (Figure 6A,B), indicating that moxGFP2 folded efficiently into a fluorescent form in the ER lumen. Moreover, when fused to Kar2, moxGFP2 showed similar *in vivo* photostability and brightness as msGFP2 (Figure 6C).

**FIGURE 6:**
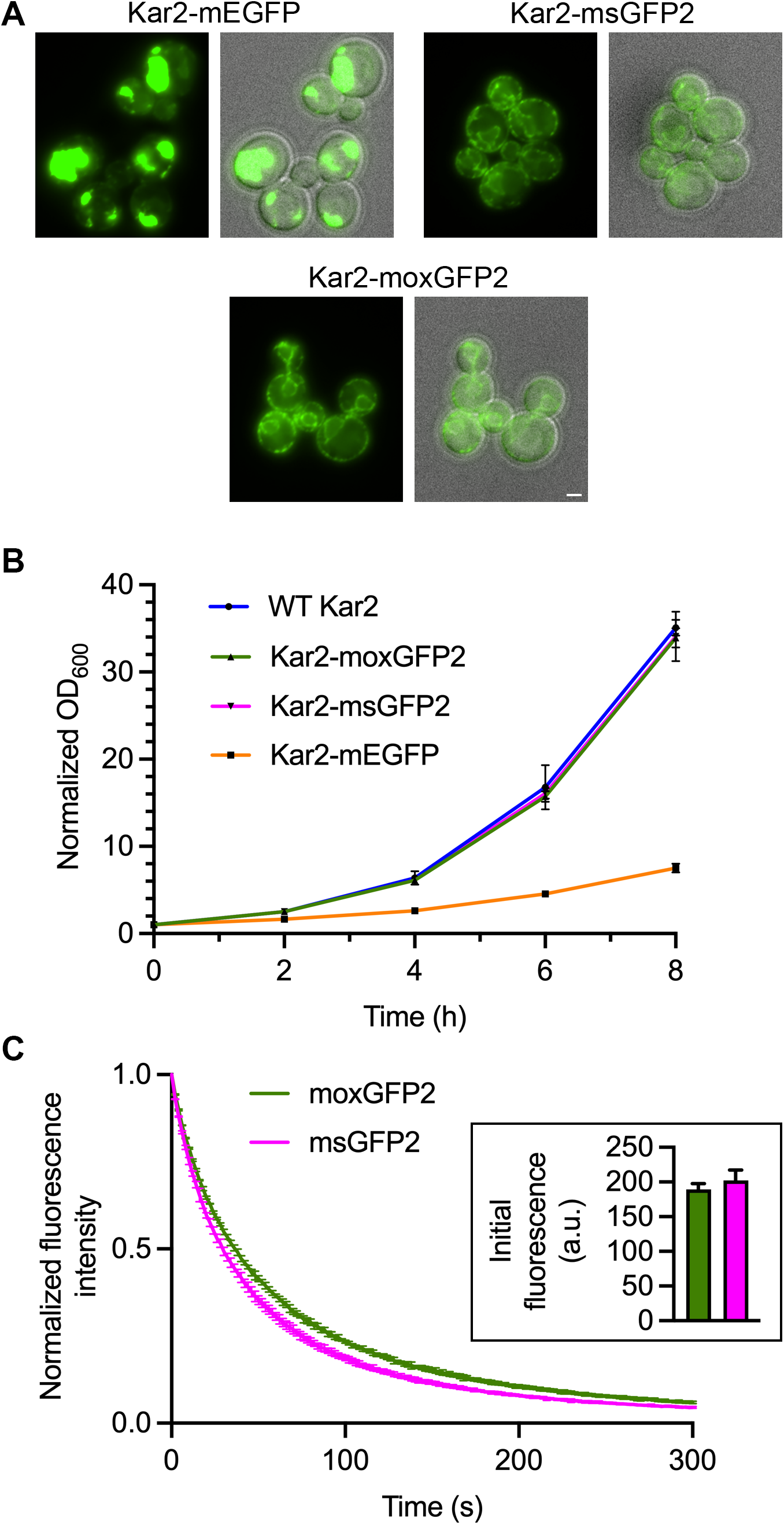
moxGFP2 is suitable for use as a fluorescent tag in the yeast ER lumen. (A) Fluorescence and combined fluorescence/brightfield images of yeast strains in which the endogenous *KAR2* gene was replaced with the indicated GFP fusion gene. Scale bar, 2 µm. (B) Growth in rich liquid medium of yeast strains expressing either wild-type (“WT”) Kar2 or the indicated GFP fusions to Kar2. The optical densities at 600 nm (OD_600_) were approximately 0.1 at the beginning of the experiment, and those initial values were normalized to 1.0. (C) Photostability of moxGFP2 and msGFP2 fused to Kar2 in live yeast cells. Confocal z-stacks were captured every 2 s for 5 min and then average projected to create movies. Plotted are averaged fluorescence signals from three movies per strain, where each movie included approximately 10-15 cells. Mean fluorescence signals were normalized to the values at time zero. The bars represent SEM values. Shown in the inset are mean cellular fluorescence signals with SEM values for the first time points of the Kar2-moxGFP2 movies (green) and the Kar2-msGFP2 movies (magenta).

As a second test of moxGFP2 in the lumen of the secretory pathway, we tagged GalNAc-T2 with msGFP2 or moxGFP2 and expressed these constructs in cultured RPE-1 cells. The labeling of Golgi structures was similar (Figure 7A), as was the photostability of the fluorescence signals (Figure 7B). The combined results confirm that moxGFP2 is well behaved as a tag for use in the lumen of endomembrane system compartments.

**FIGURE 7:**
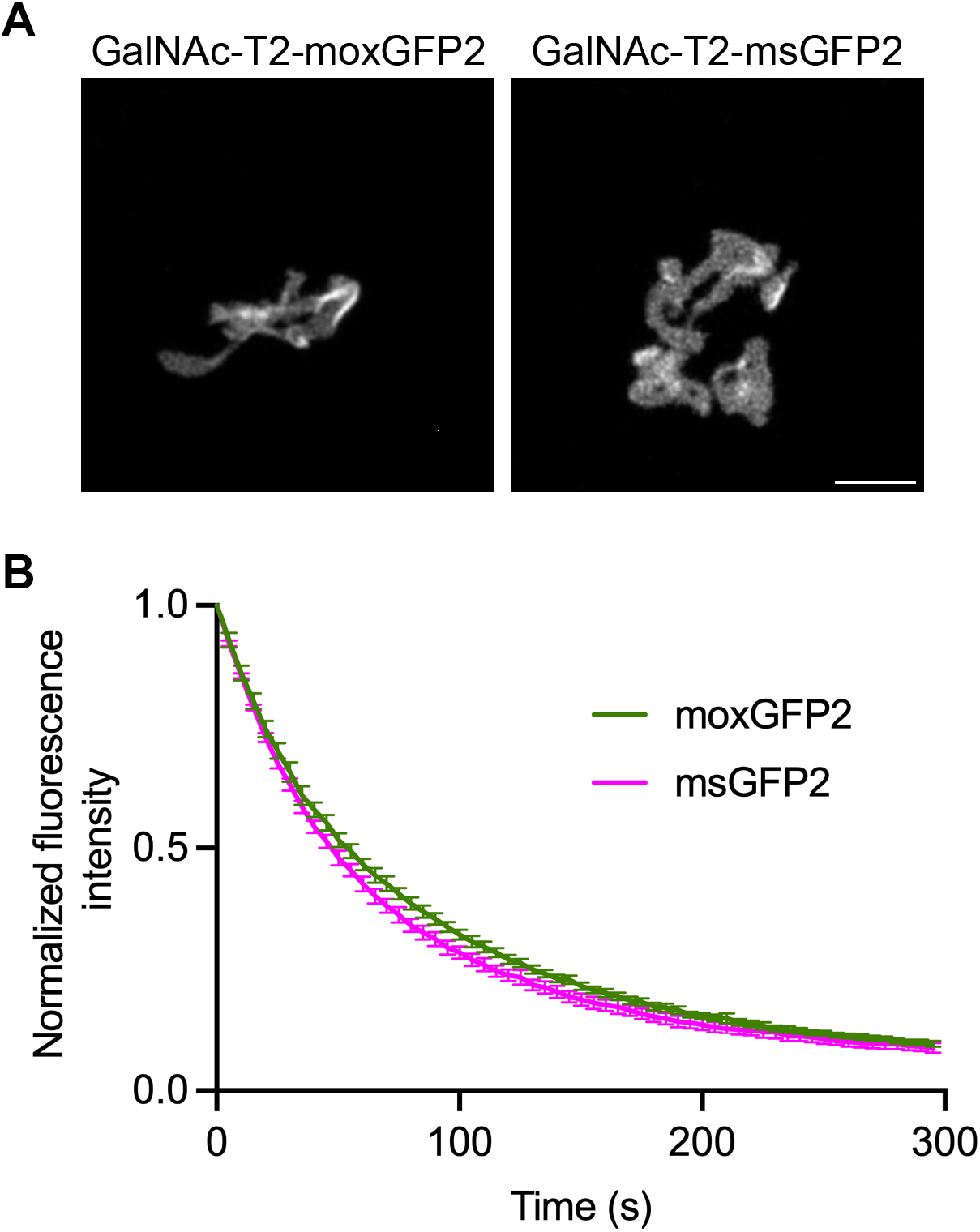
moxGFP2 is suitable for use as a fluorescent tag in the mammalian secretory pathway. (A) Labeling of the Golgi ribbon in RPE-1 cells by transient expression of constructs encoding moxGFP2 or msGFP2 fused to GalNAc-T2. Scale bar, 5 µm. (B) Photostability of moxGFP2 and msGFP2 fused to GalNAc-T2 in live RPE-1 cells. Four cells from each sample were imaged by capturing confocal z-stacks every 5 s for 5 min, and the signals from the Golgi structures were quantified. Mean fluorescence signals were normalized to the values at time zero. The bars represent SEM values.

### Conclusions

Although FPs with special characteristics are useful for certain purposes (Nienhaus and Nienhaus, 2022), the most common application of FPs is simple live-cell imaging of tagged proteins. Therefore, a worthwhile goal is to engineer monomeric FPs that fold efficiently and exhibit high brightness and photostability. Here we describe msYFP2, which is a good choice for many experimental systems that require a yellow FP. msYFP2 is less bright than the recently described mGold (Lee et al., 2020), but msYFP2 is comparable in photostability to mGold, and msYFP2 has superfolder properties that should ensure efficient folding in a variety of cellular environments. We also describe moxGFP2, which is a good choice for experimental systems that require placing tags in oxidizing environments such as the lumen of the secretory pathway. When compared to superfolder GFPs for use in oxidizing environments, moxGFP2 has two advantages: the lack of cysteines in moxGFP2 will prevent disulfide bond formation under any circumstances, and the lower folding stability of moxGFP2 should make its post-translational translocation more efficient than that of superfolder GFP (Fitzgerald and Glick, 2014). We now routinely use moxGFP2 to tag the luminal portions of yeast Golgi proteins, whereas that task was challenging with early GFP derivatives (Wooding and Pelham, 1998). It is remarkable that three decades after *A. victoria* GFP was first harnessed as a fluorescent label, this research tool is still being refined.

## MATERIALS AND METHODS

### DNA constructs

Plasmids were constructed using standard methods, including In-Fusion cloning (TaKaRa). Cloning was performed with commercially prepared Stellar competent cells (TaKaRa). Key plasmids were fully sequenced using plasmidsaurus and were deposited with Addgene, accompanied by annotated SnapGene files.

### Bacterial cell procedures

Assays involving bacterial expression were performed using XL1-Blue competent cells. Expression and purification of 6xHis-tagged FPs were performed as previously described (Valbuena et al., 2020). Superfolder activity assays with proinsulin-FP fusions expressed in bacteria, and immunoblotting of whole bacterial cell extracts using an anti-GFP antibody, were performed as previously described (Valbuena et al., 2020).

### Yeast cell procedures

The parental *S. cerevisiae* strain was JK9-3da (Kunz et al., 1993). Tagging of proteins with C-terminal FPs was performed by linearizing the plasmids for pop-in/pop-out gene replacement (Rothstein, 1991; Rossanese et al., 1999) using BsiWI for *SEC7* or XbaI for *KAR2*. In the case of *KAR2*, the FP tags were inserted upstream of the C-terminal HDEL sequence.

Prior to imaging, cells were grown in nonfluorescent minimal glucose medium (Bevis et al., 2002) in baffled flasks with shaking. The growth temperature was 23°C for strains with tagged Sec7 or 30°C for strains with tagged Kar2. 4D imaging of cells immobilized on coverglass-bottom dishes was performed with a Leica Stellaris confocal microscope using a Plan-Apo 1.4-NA 63x oil immersion objective, with z-stacks of 22 0.3-µm optical slices collected every 2 s (Johnson and Glick, 2019). Deconvolution of the images and quantification of the fluorescence signals were carried out as previously described (Johnson and Glick, 2019). Widefield imaging of cells compressed on a slide beneath a coverslip was performed by capturing a single image in a plane near the centers of the cells using a Zeiss Axio Observer microscope with a Plan-Apo 1.4-NA 63x oil immersion objective.

To measure growth curves, the cells were grown in rich glucose medium in baffled flasks with shaking at 30°C. After growth overnight to mid-log phase, the cultures were diluted and grown at 30°C for an additional 8 h.

### Mammalian cell procedures

hTERT-immortalized retinal pigment epithelial (RPE-1) cells were obtained from the American Type Culture Collection (ATCC). The cells were maintained in DMEM/F12 media (ATCC) supplemented with 10% FBS (ATCC) and 0.01 mg/ml Hygromycin B (Invitrogen) at 37°C and 5% CO_2_.

Transient transfection with DNA constructs was performed as follows. The cells were trypsinized with TrypLE (Gibco) for 10-15 min at 37°C, centrifuged at 100 rcf for 3.5 min, and washed once with 1 ml of PBS (Gibco). After analysis with an automated cell counter (Nexcelom), ∼250,000 cells were centrifuged at 100 rcf for 3 min and resuspended in 12 µl of electroporation buffer (Invitrogen). 50-100 ng of plasmid DNA were added to the cell suspension. The cells were then loaded into a 10 µl electroporation tip (Invitrogen) and electroporated using the Neon Transfection system (Invitrogen) with these parameters: voltage = 1350, width = 20, pulses = 3. Electroporated cells were immediately plated onto glass bottom dishes (MatTek) containing DMEM/F12 media supplemented with 20% FBS. The cells were allowed to recover at 37°C. At 24 h after electroporation, the media was changed to FluoroBrite DMEM (Gibco) supplemented with 20% FBS.

Cells were imaged 2 days after electroporation. Imaging was performed at 37°C using a Leica SP5 2-photon microscope equipped with a Plan-Apo 1.4-NA 63x oil immersion objective. 2 h prior to imaging, the culture medium was supplemented with 25 mM HEPES (Cytiva) and 100 µg/mL cycloheximide (RPI Research Products International). For fluorescence excitation, an argon laser was set to 488 nm for GFP or 514 nm for YFP. For fluorescence emission, the GFP collection window was set to 495-578 nm and the YFP collection window was set to 520-603 nm. Individual Golgi ribbons were imaged using a zoom value of 10, pinhole value of 1 AU, line average of 2, and z-step size of 170 nm. Each frame was 512 x 512 pixels of size 48 nm x 48 nm. Full z-stacks of the Golgi ribbon (40-50 optical sections) were collected every 5 s for 5 min. The z-stacks were deconvolved and then average projected.

### Drosophila procedures

*Drosophila yki* was endogenously tagged with msYFP2 via CRISPR/Cas9 using a linker sequence identical to the one used for the previously generated Yki-Venus (Manning et al., 2018). To generate the guide RNA (gRNA), complementary oligonucleotides encoding a guide sequence at the 3’ end of the *yki* coding sequence, GACAAACCTGACGATTTGGAA, were annealed, phosphorylated using polynucleotide kinase, and ligated into BbsI-cut pU6-BbsI-chiRNA, which is designed to express the gRNA under control of the *Drosophila* U6 promoter (Gratz et al., 2013). To generate a donor template, the msYFP2 coding sequence preceded by a linker (5’-CCTACGCCCCCAACTGAGAGAACTCAAAGGTTACCCCAGTTGGGGCACTACGGCGCGCCCAA-3’ encoding the amino acid sequence PTPPTERTQRLPQLGHYGAPK) was flanked on the 5’ end by 962 bp of *yki* genomic sequence upstream of the stop codon (to generate a 5’ homology region, or 5’HR) and on the 3’ end by the 950 bp of *yki* genomic sequence immediately downstream of the stop codon (3’HR). The resultant 5’HR-linker-msYFP2-3’HR sequence was cloned into the pBluescript II SK(+) vector. For injection, the donor template along with the gRNA were sent to a third-party facility (GenetiVision Corporation). Putative transformants were pre-screened by examining green fluorescence in third instar larvae, followed by PCR screening and sequencing to ensure proper integration of the donor sequence.

Photostability tests were performed with wing imaginal tissue explants from *yki-msYFP2/+* or *yki-Venus/+* wandering third instar larvae. Tissues were dissected in Schneider’s Drosophila Medium (Sigma) supplemented with 10% Fetal Bovine Serum (Thermo Fisher Scientific) on a siliconized glass slide. The explants were then transferred with a pipette in 5-10 µl of medium to a glass bottom microwell dish (MatTek, 35 mm petri dish, 14 mm microwell) with No. 1.5 coverglass. The tissues were oriented so that the apical side of the disc proper faced the coverglass. A Millicell culture insert (Sigma, 12 mm diameter, 8 µm membrane pore size) was prepared in advance by cutting off the bottom legs with a razor blade and removing any excess membrane material around the rim of the insert. The insert was carefully placed into the 14 mm microwell space, with the membrane directly overlaying the tissues. The space between the insert and the microwell was sealed with ∼15 µl of mineral oil, and 200 µl of dissection medium was added into the insert chamber. Tissues of both genotypes were imaged under identical conditions on a Zeiss LSM 880 laser scanning confocal microscope equipped with a 514 nm argon laser, GaAsP spectral detector, Plan-Apo 1.4-NA 40x oil immersion objective, and Zen Black software. Five tissues per genotype were scanned, and all data were acquired from the anterior ventral region of the wing imaginal discs. z-stacks of 4 slices (2.9 µm deep from the apical surface of the wing disc proper) with an area of ∼35 x 35 µm were acquired continuously in bidirectional mode for a total of 20 scans with 4x averaging and a scan speed of 8. Maximum intensity projections of the time-lapse images were generated in Fiji (ImageJ), and fluorescence intensities were measured using the Plot Z-axis Profile function. The measurements were background subtracted (using background fluorescence measurements of nonfluorescent tissues from *w^1118^* larvae) and normalized to the maximum intensity values.

To examine Yki-msYFP2 nuclear localization, the following cross was performed:

*Yki-msYFP2/CyO, dfdYFP X wts RNAi; hh>Gal4, UAS-mChNLS/TM6, Tb, tub>Gal80*

Non-balancer third instar larvae were selected, and wing imaginal discs were dissected and mounted as described above. A single optical section through the basal plane of the tissue (where the nuclei are clearly identifiable) was acquired sequentially using lasers of wavelength 514 nm (for YFP) and 561 nm (for mCherry).

### Measurements of FP properties

Photobleaching assays with purified FPs were performed as previously described (Valbuena et al., 2020) except that a Zeiss Axio Observer widefield microscope was used.

Excitation and emission spectra were measured with a Duetta 3-in-1 spectrofluorometer (HORIBA Scientific). Dilution series for each FP were generated based on the absorbance at 280 nm in PBS buffer, and excitation and emission spectra were measured for each dilution in quartz cuvettes. For GFP variants, excitation was measured from 450 to 600 nm (emission wavelength 510 nm, 1 nm excitation step increment) and emission was measured from 500 to 650 nm (excitation wavelength 488 nm). For YFP variants, absorbance was measured from 450 to 600 nm (emission wavelength 527 nm, 1 nm excitation step increment) and emission was measured from 520 to 700 nm (excitation wavelength 513 nm). Excitation and emission bandpasses were set to 3 nm for all samples. The built-in instrument software was used to calculate the integrated fluorescence emission for each sample. Linear plots of integrated fluorescence versus absorbance at 280 nm were compared to those of EGFP and EYFP to determine quantum yields (Valbuena et al., 2020).

### Software analysis

DNA constructs were designed and analyzed using SnapGene (Dotmatics). Numerical data were plotted and analyzed using Prism (Dotmatics). Figures were assembled with ImageJ or Fiji together with Photoshop (Adobe). Deconvolution was performed using Huygens Essential (Scientific Volume Imaging).

## ACKNOWLEDGMENTS

This work was supported by NIH grants R35 GM144050 to BSG and R01 NS034783 to RGF. FMV, AHK, SAT, and ACG were supported by NIH training grant T32 GM007183. FMV was also supported by an HHMI Gilliam Felllowship, and SAT was also supported by an NSF Graduate Research Fellowship. For assistance with fluorescence microscopy, we thank Christine Labno at the Integrated Microscopy Core Facility, which is supported by the NIH-funded Cancer Center Support Grant P30 CA014599. Thanks for assistance with the collection of spectra to Elena Solomaha at the Biophysics Core Facility and Justin Jureller at Chicago Materials Research Center, and thanks for assistance with screening CRISPR fly lines to Nicki Nouri. Additional thanks to the Glick lab for constructive feedback.

---

**FIGURE S1:**
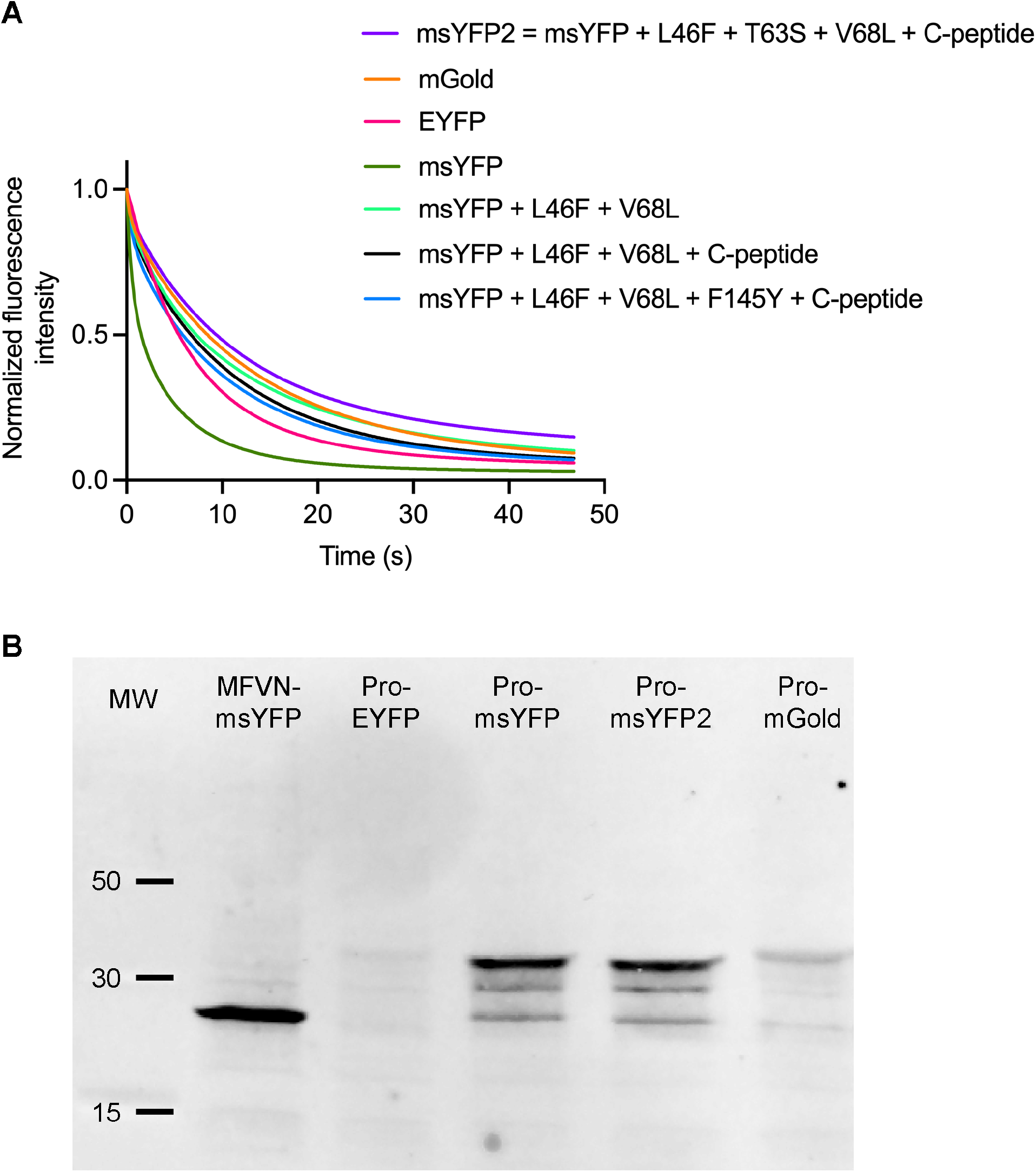
Supporting data for Figure 1. (A) Photobleaching of purified YFP variants. The indicated YFPs were imaged repeatedly by widefield microscopy as in Figure 1A. “C-peptide” indicates replacement of the original msYFP C-terminal peptide with the C-terminal peptide from msGFP2. (B) Expression levels of YFP variants in *E. coli*. The indicated YFPs were expressed as fusions to proinsulin (“Pro”), and whole cell extracts were separated by SDS-PAGE followed by immunoblotting with an anti-GFP antibody. The sample labeled “MFVN-msYFP” was msYFP preceded by the first four codons of proinsulin. “MW” is molecular weight markers, with the sizes indicated in kDa.

**FIGURE S2:**
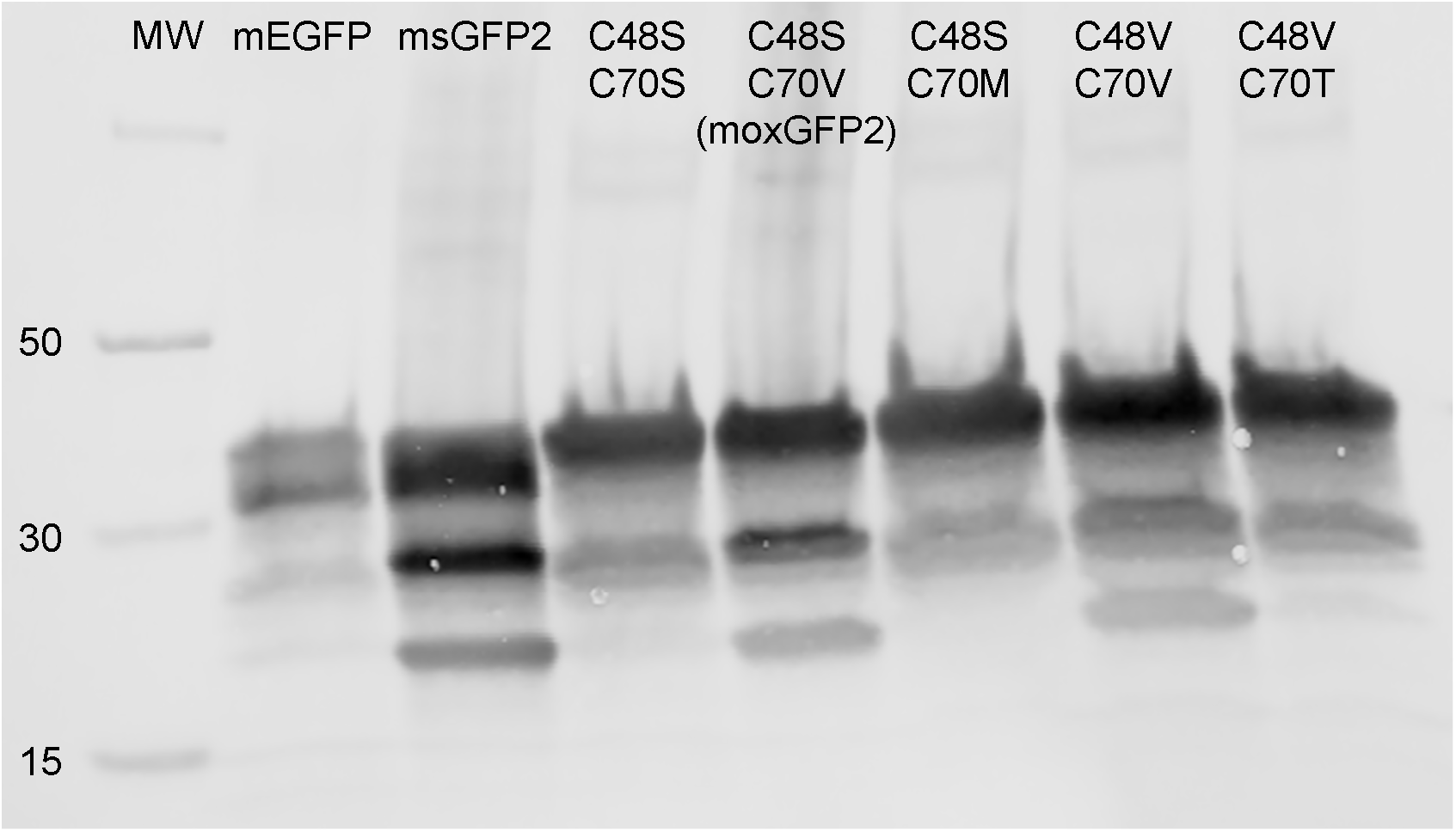
Expression levels of mEGFP and of msGFP2 derivatives in *E. coli* (supporting data for Figure 5A). The indicated GFPs were expressed as fusions to proinsulin, and whole cell extracts were separated by SDS-PAGE followed by immunoblotting with an anti-GFP antibody. Cysteine substitutions indicate mutant derivatives of msGFP2. “MW” is molecular weight markers, with the sizes indicated in kDa.

## Abbreviations used

FP: fluorescent protein
GalNAc-T2: N-acetylgalactosaminyltransferase 2
GFP: green fluorescent protein
YFP: yellow fluorescent protein
Yki: Yorkie
Wts: Warts

